# Characterization of the complete mitochondrial genome of the New Zealand parasitic blowfly *Calliphora vicina* (Insecta: Diptera: Calliphoridae)

**DOI:** 10.1101/2020.11.05.370361

**Authors:** Nikola Palevich, Luis Carvalho, Paul Maclean

## Abstract

In the present study, the complete mitochondrial genome of the New Zealand parasitic blowfly *Calliphora vicina* (blue bottle blowfly) field strain NZ_CalVic_NP was generated using next-generation sequencing technology and annotated. The 16,518 bp mitochondrial genome consists of 13 protein-coding genes, two ribosomal RNAs, 22 transfer RNAs, and a 1,689 bp non-coding region, similar to most metazoan mitochondrial genomes. Phylogenetic analysis showed that *C. vicina* NZ_CalVic_NP does not form a monophyletic cluster with the remaining three Calliphorinae species. The complete mitochondrial genome sequence of *C. vicina* NZ_CalVic_NP is a resource to facilitate future species identification research within the Calliphoridae.

---

The diminished efficacy demonstrated by current members of the Calliphoridae (blowflies) treatments due to the emergence of resistance in blowflies against many classes of insecticides calls for improved DNA-based diagnostics tools. High-level phylogenetic relationships within the Calliphoridae are still largely unresolved primarily due to their large and highly variable mitochondrial (mt) genomes of blowflies. *Calliphora vicina* NZ_CalVic_NP was selected for genome sequencing as a representative of an NZ field strain of *C. vicina*.

The *C. vicina* specimen was collected from the Palmerston North area (40°21.3′ S, 175°36.7′ E), and is stored and available upon request from AgResearch Ltd., Grasslands Research Centre (accession number: NPY120886). High molecular weight genomic DNA was isolated from entire *C. vicina* adult males using a modified phenol:chloroform protocol explained in our previous articles (Palevich et al. 2019a; Palevich et al. 2019b; Palevich et al. 2019d). The Illumina NovaSeq™ 6000 (PE150, Novogene, China) platform was used to amplify the entire mitochondrial genome sequence. The mitochondrial genome was assembled and annotated as previously described (Palevich et al. 2019c; Palevich et al. 2019e; Palevich et al. 2020).

The length of complete mitochondrial genome is 16,518 bp, with the overall 77.8% AT content (BioProject ID: PRJNA667961, GenBank accession number: MW123003). The overall nucleotide distribution for the mitochondrial genome is 39.5 % A, 13.0 % C, 9.2 % G, and 38.1 % T. The structure of the mitochondrial genome is typical of insect mitochondrial genomes (Cameron 2014) which consists of 13 protein-coding genes, 22 transfer RNAs, and 2 ribosomal RNAs. Among these 37 genes, 23 genes encoded on the majority strand while remaining 14 genes encoded on the minority strand. There are eight more complete mitochondrial genomes recorded belong to the genus Calliphora (*C. vicina, C. vomitoria, C. nigribarbis* and *C. chinghaiensis*) (Nelson et al. 2012; Chen et al. 2016; Ren et al. 2016; Karagozlu et al. 2019). In comparison, the reported *C. vicina* NZ_CalVic_NP has the longest complete mitochondrial genome and the size difference with the shortest record is 1,249 bp (*C. chinghaiensis*). The main reason for the size difference is the control region. The entire ‘control region’ that is non-coding and AT-rich lies between the 12*S* rRNA and tRNA-Ile in insect mitochondrial genomes, and this area in the *C. vicina* NZ_CalVic_NP is 1,689 bp in length which is the longest among all Calliphora records.

The phylogenetic position of *C. vicina* NZ_CalVic_NP within the family Calliphorinae was estimated using maximum-likelihood, implemented in RAxML version 8.2.11 (Stamatakis 2014), and the Bayesian inference (BI), implemented in MrBayes version 3.2.6 (Huelsenbeck et al. 2001) approaches using default settings.

For analysis, the phylogenetic tree was reconstructed using the complete mitogenome sequences of available blowfly species and isolates retrieved from GenBank with the 13 concatenated mitochondrial PCGs and rRNA genes (Figure 1). *Calliphora vomitoria* was the most related species with *C. nigribarbis* and *C. chinghaiensis*. Overall, the phylogenetic topology is similar to previous studies (Chen et al. 2016), suggesting that the genus Calliphora is not monophyletic. This study provides additional complete mitogenome data for the improvement and future investigation of the Calliphoridae phylogeny.

**Figure 1.**
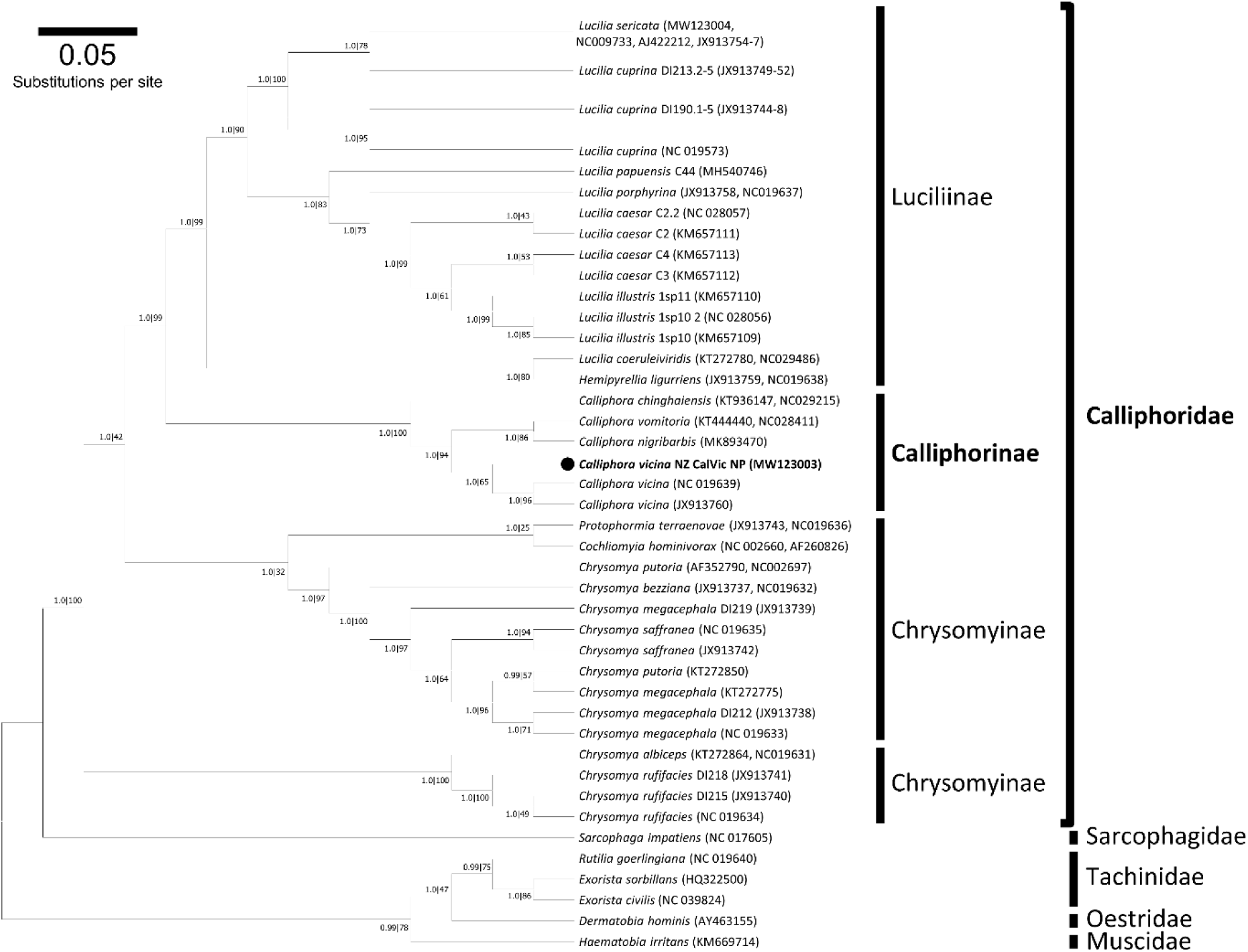
A summary of the molecular phylogeny of the Calliphoridae complete mitochondrial genomes. The evolutionary relationship of *C. vicina* field strain NZ_CalVic_NP (black circle) was compared to the complete mitochondrial genomes of 68 blowfly species or isolates retrieved from GenBank (accession numbers in parentheses) and nucleotide sequences of all protein-coding genes were used for analysis. Phylogenetic analysis was conducted using the Bayesian approach implemented in MrBayes version 3.2.6 (Huelsenbeck et al. 2001) and maximum likelihood (ML) using RAxML version 8.2.11 (Stamatakis 2014). The mtREV with Freqs. (+F) model was used for amino acid substitution and four independent runs were performed for 10 million generations and sampled every 1,000 generations. For reconstruction, the first 25% of the sample was discarded as burnin and visualized using Geneious Prime (Kearse et al. 2012). Nodal support is given: Bayes posterior probabilities|RAxML bootstrap percentage. The phylogram provided is presented to scale (scale bar = 0.05 estimated number of substitutions per site) with the species *Haematobia irritans* from the family Muscidae used as the outgroup.

## Data availability statement

The data that support the findings of this study are openly available in GenBank of NCBI at https://www.ncbi.nlm.nih.gov, reference number MW123003.

